# Reduction of ribosomal expansion segments in yeast species of the *Magnusiomyces/Saprochaete* clade

**DOI:** 10.1101/2023.07.14.548829

**Authors:** Filip Brázdovič, Broňa Brejová, Barbara Siváková, Peter Baráth, Tomáš Vinař, Ľubomír Tomáška, Jozef Nosek

**Affiliations:** Faculty of Natural Sciences, Comenius University in Bratislava, Slovakia; Faculty of Mathematics, Physics, and Informatics, Comenius University in Bratislava, Slovakia; Institute of Chemistry, Slovak Academy of Sciences, Bratislava, Slovakia

**Keywords:** expansion segments, ribosome, ribosomal RNA, Magnusiomyces, yeast

## Abstract

Ribosomes are ribonucleoprotein complexes highly conserved across all domains of life. The size differences of ribosomal RNAs (rRNAs) can be mainly attributed to variable regions termed expansion segments (ESs) protruding out from the ribosomal surface. The ESs were found to be involved in a range of processes including ribosome biogenesis and maturation, translation, and co-translational protein modification. Here, we analyze the rRNAs of the yeasts from the *Magnusiomyces/Saprochaete* clade belonging to the basal lineages of the subphylum Saccharomycotina. We find that these yeasts are missing more than 400 nt from the 25S rRNA and 150 nt from the 18S rRNAs when compared to their canonical counterparts in *Saccharomyces cerevisiae*. The missing regions mostly map to ESs, thus representing a shift toward a minimal rRNA structure. Despite the structural changes in rRNAs, we did not identify dramatic alterations of the ribosomal protein inventories. We also show that the size-reduced rRNAs are not limited to the species of the *Magnusiomyces/Saprochaete* clade, indicating that the shortening of ESs happened independently in several other lineages of the subphylum Saccharomycotina.

**Significance:** Expansion segments are variable regions present in the ribosomal RNAs involved in the ribosome biogenesis and translation. Although some of them were shown to be essential, their functions and the evolutionary trajectories leading to their expansion and/or reduction are not fully understood. Here, we show that the yeasts from the *Magnusiomyces/Saprochaete* clade have truncated expansion segments, yet the protein inventories of their ribosomes do not radically differ from the species possessing canonical ribosomal RNAs. We also show that the loss of expansion segments occurred independently in several phylogenetic lineages of yeasts pointing out their dispensable nature. The differences identified in yeast ribosomal RNAs open a venue for further studies of these enigmatic elements of the eukaryotic ribosome.

## Introduction

Ribosomes are essential molecular machines carrying out the translation of genetic information from messenger RNA molecules into amino acid sequences of polypeptide chains. Although the core of these nucleoprotein complexes is highly conserved, many changes in both RNA and protein components occurred since the divergence from the last common ancestor of all living organisms (Bowman et al. 2020). This can be exemplified by differences between bacterial and eukaryotic ribosomes. While the 70S ribosome from bacteria *Escherichia coli* contains 3 ribosomal RNA (rRNA) molecules (23S, 16S, 5S), 54 proteins and reaches the size of 2.3 MDa, its 80S counterparts from baker’s yeast *Saccharomyces cerevisiae* and human are composed of four rRNAs (25S/28S, 18S, 5.8S, 5S) and 79-80 proteins and reach sizes of 3.3 MDa and 4.3 MDa, respectively (Melnikov et al. 2012). During the evolution, ribosome has increased its size in several phases by means of accretion (Bokov and Steinberg 2009; Petrov et al. 2014a; Petrov et al. 2015; Biesiada et al. 2022), the most recent phase being the emergence of expansion segments (ESs) and their further enlargement in higher eukaryotes (Biesiada et al. 2022). The ESs are located on the surface of the ribosome not affecting its functional core and exhibit remarkable variability among different species. The enlargement of rRNA sequences due to ESs introduces a substantial burden for the cell. For example, the genome of *S. cerevisiae* contains ∼150-200 rRNA gene copies (Johnston et al. 1997), and around 60 % of all transcription in rapidly growing yeast cells is dedicated to rRNAs, enabling the production of ∼2,000 ribosomes per minute (Warner 1999). While the molecular functions of ESs still remain elusive, several studies point to their roles in ribosome biogenesis, translational control, interactions with proteins and RNA molecules, and cellular response to oxidative stress (Bradatsch et al. 2012; Pánek et al. 2013; Ramesh and Woolford 2016; Gómez Ramos et al. 2016; Shedlovskiy et al. 2017; Parker et al. 2018; Fuji et al. 2018, Mestre-Fos et al. 2019; Knorr et al. 2019,2023; Shankar et al. 2020; Leppek et al. 2020,2021; Wild et al. 2020; Krauer et al. 2021; Vos and Kothe 2022).

In this study, we perform a comparative analysis of rRNAs from arthroconidial yeast species classified to the genus *Magnusiomyces* (and its anamorph *Saprochaete*) belonging to *Dipodascaceae* family (Saccharomycotina, Ascomycota) (Supplementary Table 1). A common feature of these yeasts is atypical 18S rRNA that lacks several regions originating from ribosomal helices and ESs making it ∼150 nt shorter than canonical yeast 18S rRNA (Ueda-Nishimura and Mikata 2000). These alterations make *Magnusiomyces*/*Saprochaete* clade distinct from the sister lineages, which comprise the genera *Dipodascus* and *Galactomyces* including their anamorphic form *Geotrichum* (Ueda-Nishimura and Mikata 2000; de Hoog and Smith 2004). By comparison to the model structures of *S. cerevisiae* rRNAs (Ben-Shem et al. 2011; Petrov et al. 2014b), we show that the changes in magnusiomycete ribosomes are not limited to 18S rRNA. Several helices and ESs are missing also in 25S and 5.8S rRNAs, considerably reducing their sizes. The altered regions map to the ribosomal surface, likely affecting interactions of the ribosome with its macromolecular partners. To complement the comparative analysis of rRNAs, we also investigate ribosomal protein inventory of pathogenic yeast *Magnusiomyces capitatus* by means of both bioinformatic analysis of the nuclear genome sequence (Brejová et al. 2019a) and mass-spectrometry analysis of the proteins present in purified ribosomal fractions. Our results indicate that the genome of *M. capitatus* encodes a standard set of ribosomal proteins implying that the alterations of ESs in the magnusiomycete ribosomes have not been accompanied by changes in the ribosomal protein inventory. Further comparative analysis of rRNA sequences from other yeast species show that albeit rare, the changes of ESs occur in several additional lineages of the subphylum Saccharomycotina, illustrating the dispensable nature of these rRNA segments.

## Results and Discussion

### Ribosomal RNA genes in the species of *Magnusiomyces/Saprochaete* clade

We identified the genes for cytosolic rRNAs in the genome sequences of 16 species classified into *Magnusiomycete/Saprochaete* clade and two representatives from its sister clades *Dipodascus* (*D. albidus*) and *Galactomyces* (*G. candidus*) (Supplementary Table 1). In all examined species, these genes are organized in a cluster comprising two transcriptional units for (i) 18S, 5.8S, and 25S rRNAs, and (ii) 5S rRNA, although in some species, the gene for 5S rRNA has inverted orientation (Figure 1). The rDNA cluster is tandemly repeated and it has been estimated that the genomes of *M. capitatus* and *M. ingens* contain about 92 and 144 copies, respectively (Brejová et al. 2019a). This arrangement is typical for most Saccharomycotina species, although some yeasts classified into the *Dipodascaceae* family have a different organization of rRNA genes. For example, *Yarrowia lipolytica* possesses multiple rDNA units located in subtelomeric chromosomal regions and its rDNA cluster lacks the 5S rRNA gene whose copies are dispersed throughout the genome (Casaregola et al. 2000). Phylogenetic analysis based on the sequences of all four rRNAs indicates that, except for *S. psychrophila*, the remaining *Magnusiomyces* and *Saprochaete* species form a distinct lineage comprised of three branches: (i) *M. capitatus*, *M. clavatus*, *M. spicifer*; (ii) *M. starmeri, M. ingens, S. ingens, S. chiloensis, M. ovetensis, S. quercus* and; (iii) *S. saccharophila, S. fungicola, M. tetraspermus, S. suaveolens, S. gigas, M. magnusii* (Figure 1). In contrast to previous classification (de Hoog and Smith, 2011b), *S. psychrophila* represents a distinct lineage parallel to the genera *Dipodascus* and *Galactomyces,* indicating that this species does not belong to *Magnusiomyces*/*Saprochaete* clade. This conclusion is also in line with the results of comparative analysis of rRNAs (see below), which shows that the absence of several ESs in rRNAs clearly separates the species of *Magnusiomyces/Saprochaete* clade from the yeasts belonging to the sister lineages including *S. psychrophila*.

**Fig. 1.**
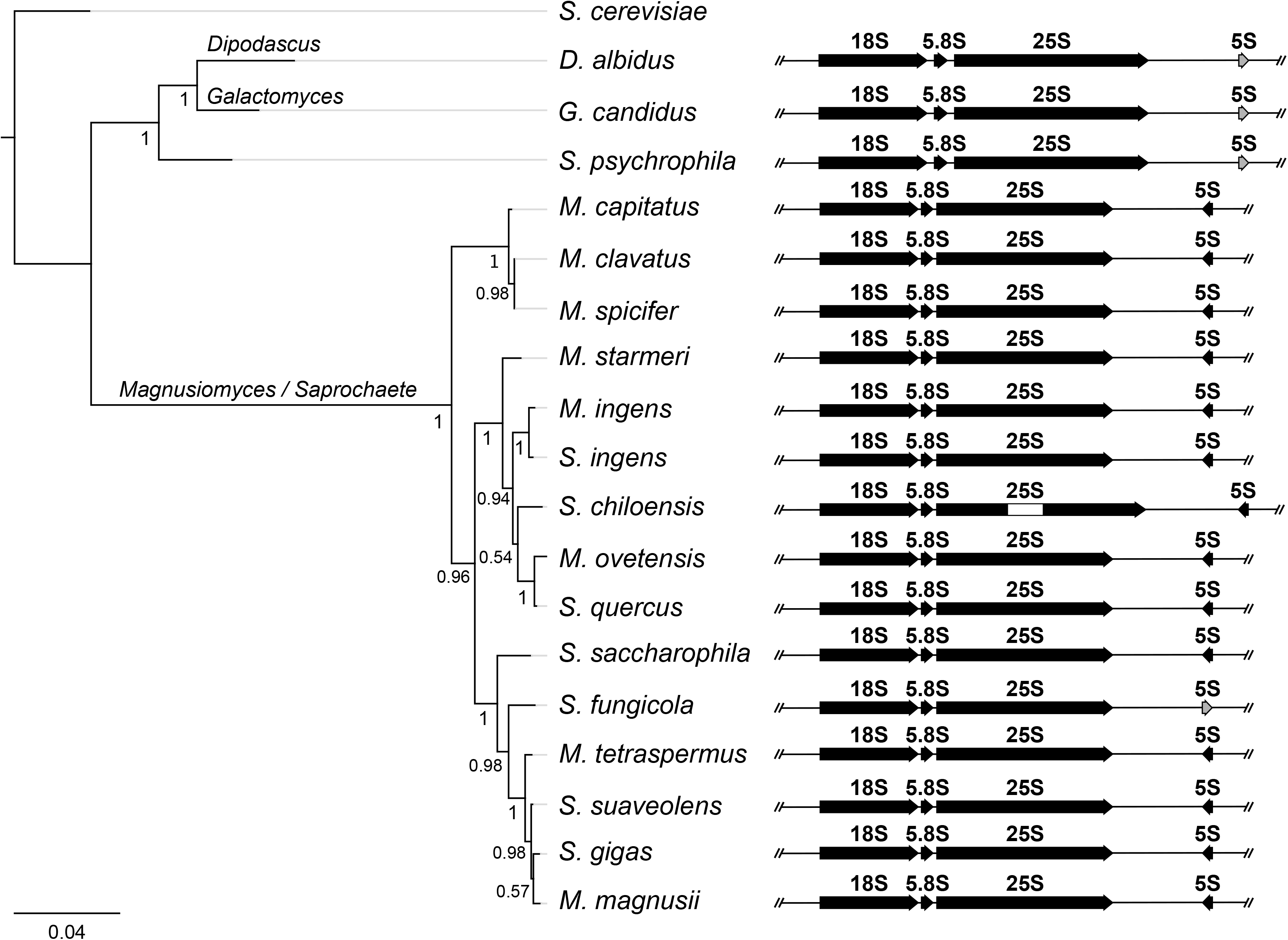
Phylogenetic tree of the *Magnusiomyces/Saprochaete* clade. The tree was built using Fasttree from concatenated alignment of rRNA sequences (see Materials and Methods). *S. cerevisiae* was used as an outgroup. Arrangement of rRNA genes in the rDNA cluster is shown for each species examined in this study (Supplementary Table 1). Note that the 5S rRNA gene in *D. albidus*, *S. psychrophila*, and *S. fungicola* has inverted orientation (shown in gray) compared to the remaining species. Moreover, the 25S rRNA gene of *S. chiloensis* contains a 437 nt long group I intron (shown as a white rectangle), and its secondary structure displays typical features of the subgroup IC1 (Supplementary Figure 1).

### Magnusiomycete rRNAs have reduced ESs

To highlight the alterations in magnusiomycete rRNAs we compared their sequences with the counterparts from *S. cerevisiae* and mapped them onto its ribosomal structure inferred from X-ray crystallography (Ben-Shem et al. 2011; Bernier et al. 2014). While 5S rRNA is highly conserved in all examined species (Supplementary Figure 2A), we identified striking alterations in the remaining three magnusiomycete rRNAs when compared to *S. cerevisiae* or closely related species *D. albidus, G. candidus*, and *S. psychrophila.* As previously described by Ueda-Nishimura and Mikata (2000), magnusiomycete 18S rRNA is shorter by ∼150 nt and lacks the ESs 9es3a, 9es3b, substantial parts of 21es6a-d, 41es10 and helices 10, 25, and 39 (numbering according to Bernier et al. 2014; Petrov et al. 2014b, Figure 2A, Supplementary Figure 2B, Supplementary Figure 3A). However, the alterations are not limited to 18S rRNA. The partial absence of helix 7 and 9ES3 makes the 5.8S rRNA molecule ∼15 nt shorter than its *S. cerevisiae* ortholog (Supplementary Figure 2C, Supplementary Figure 3B). Perhaps most striking are differences observed in 25S rRNA, which is more than 400 nt shorter than its *S. cerevisiae* counterpart due to the absence of several helices and ESs (Figure 2B, Supplementary Figure 2D, Supplementary Figure 3B). The most prominent differences occur in helices 25ES7a-c, 30, 31ES9, 63ES27a-b, 79ES31a-b, 98 and its expansion segment 98ES39b (Figure 2B, Supplementary Figure 2D, Supplementary Figure 3B). When projected on the 3D structure of *S. cerevisiae* cytosolic ribosome, the majority of these alterations occur on its surface (Figure 3, Supplementary Movie 1).

**Fig. 2.**
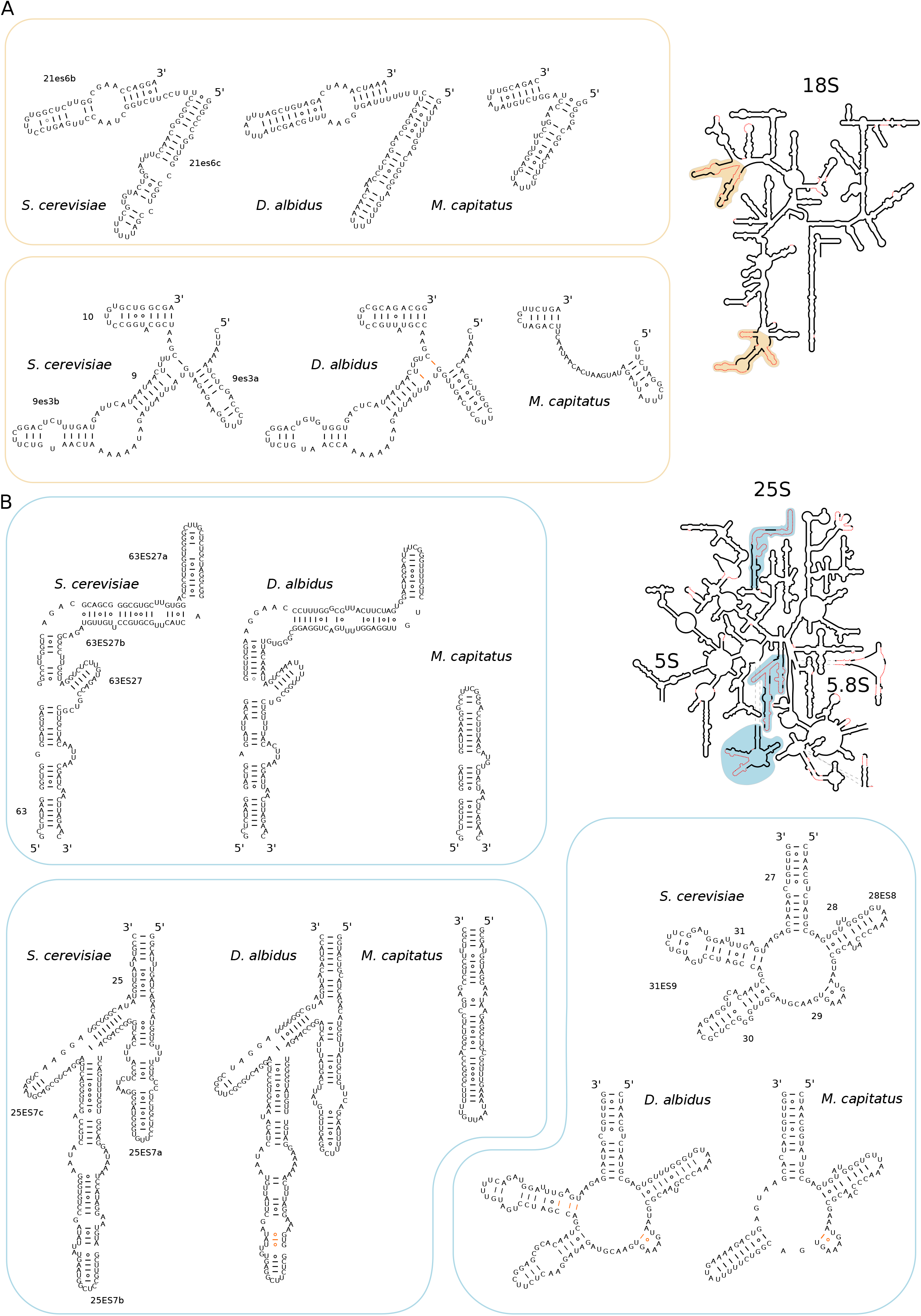
Secondary rRNA structure comparison. Shown are rRNA segments from *S. cerevisiae* (Petrov et al. 2014b), *M. capitatus,* a representative species of yeasts from *Magnusiomyces/Saprochaete* clade with reduced rRNAs, and *D. albicans,* a yeast with canonical rRNAs phylogenetically close to magnusiomycetes. The secondary structure predictions of rRNA regions from *M. capitatus* and *D. albicans* were created with the MFold program (Zuker 2003). Where needed, the structures were adjusted manually, indicated by orange color. The bonds between nucleotides are indicated with solid lines or, in the case of G○U wobble base pairs, with empty circles. The helix labels are shown in *S. cerevisiae* structures. The figure shows rRNA regions of 18S (**A**) involving helices 9 – 10 and expansion segments of helix 21, 21es6b and 21es6c and rRNA of 25S (**B**) involving helix 25 and its expansion segments 25ES7a-c, helices 27, 28, 29, 30, 31, and its expansion segment 31ES9, and region containing helix 63 and its expansion segments 63ES27, 63ES27a-b. The full SSU and LSU structures shown in the right part of the figure show the segments missing in *M. capitatus* by dotted red lines. The full-scale figures of SSU and LSU structures can be found in Supplementary Figure 3A and 3B.

**Fig. 3.**
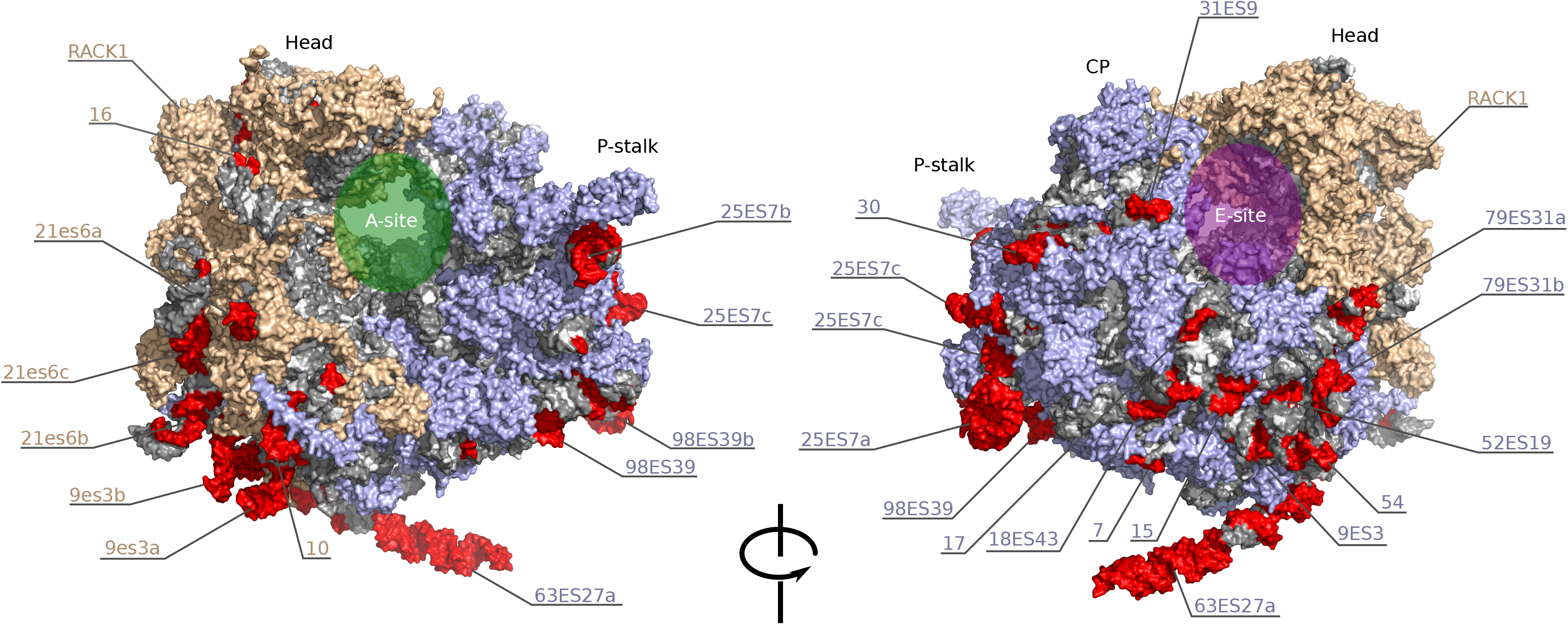
The 3D structure of the cytosolic ribosome illustrating the changes specific to the yeasts from the *Magnusiomyces/Saprochaete* clade. *S. cerevisiae* cytosolic ribosome (PDB: 4V88) (Ben-Shem et al. 2011) was used as a template. rRNA molecules are in gray, missing helices and rRNA segments are in red. Light brown color was used to indicate proteins and labels of rRNA segments of SSU. Light blue color was used to indicate proteins and labels of rRNA segments of LSU. P-stalk, central protuberance (CP) Head, and RACK1 protein were used as landmarks. Structures of helices 25ES7a and 63ES27a of 25S rRNA are taken from Knorr et al. (2019) (PDB: 6HD7).

While the sequences and structures of rRNAs from *D. albidus*, *G. candidus*, and *S. psychrophila* are similar to those of *S. cerevisiae,* the reduction of ESs in the species of *Magnusiomyces/Saprochaete* clade likely caused substantial structural changes. Predictions by Mfold (Zucker, 2003) show that the absence of helices 9es3a, 9es3b, and 10 of 18S rRNA results in a formation of two short stem loops connected by an unstructured stretch of RNA composed of the remnants of the helix 9 (Figure 2A, Supplementary Figure 2B). The segments of 18S rRNA containing helices 21ES6c and 21ES6b are substantially shorter but the overall shape seems to be retained (Figure 2A, Supplementary Figure 2B). On the other hand, the branched structures of helices 25ES7a-c, 30, 31ES9, and 63ES27a-b of 25S are reduced to single-stem loops of smaller sizes (Figure 2B, Supplementary Figure 2D). Even though these ESs were not lost completely in *Magnusiomyces/Saprochaete* clade and it is conceivable that their remnants may still serve in some interactions, the loss of their substantial portions indicates that they have likely become obsolete in magnusiomycetes.

### Magnusiomycete ribosomes are composed of a standard set of ribosomal proteins

We searched the annotated genome sequence of *M. capitatus* (Brejová et al. 2019a) for counterparts of 33 proteins of the small subunit (SSU) and 48 proteins of the large subunit (LSU) of the *S. cerevisiae* ribosome. Note that the LSU protein set includes four homologs of P1/P2 proteins. We also searched the genome sequence of *D. albidus* (Shen et al. 2018), which is phylogenetically related to *M. capitatus* but, similarly to *S. cerevisiae,* possesses canonical rRNAs. In total, we identified 81 genes encoding putative ribosomal proteins in *M. capitatus* and 107 in *D. albidus* (Supplementary Table 3). The list includes orthologs of all ribosomal proteins from *S. cerevisiae*. The ribosomal proteins of *M. capitatus* have similar sizes as their *S. cerevisiae* homologs, their sequence identity varies from 47 to 91 % and the results from RNA-Seq analysis (Brejová et al. 2019a) show that corresponding genes are highly expressed (percentile rank 97.6 – 100 %, Supplementary Table 3). Worth mentioning is the protein uL22 (MCA_06292_1) that is about 15 and 14 amino acid (AA) residues shorter compared to *S. cerevisiae* and *D. albidus* homologs, respectively, suggesting an adaptation to the reduction of ESs located in its proximity (i.e., ES7, ES27) near the ribosome exit tunnel. In *S. cerevisiae*, this protein and helix 63ES27a were recently shown to serve as universal adapter sites for N-terminal acetylation complex NatB (Knorr et al. 2023).

To complement the bioinformatic analyses, we prepared ribosomal fractions from both *M. capitatus* and *D. albidus* (Supplementary Figure 3) and investigated their protein composition by LC-MS/MS. In total, we identified 127 proteins in *M. capitatus* and 179 proteins in *D. albidus* (Supplementary Table 4). In *M. capitatus*, these included 79 ribosomal proteins (Supplementary Table 3), 30 proteins whose homologs physically associate with ribosomal proteins in *S. cerevisiae* (SGD Database, https://www.yeastgenome.org/), 9 homologs of mitochondrial ribosomal proteins, and 9 additional co-purifying proteins (Supplementary Table 4). This set contains all core ribosomal proteins, including three P1/P2 homologs, except for the homolog of eL41. Since we identified a candidate gene encoding this protein in *M. capitatus* genome, we assume that its short length (25 AAs) precluded its identification in the proteomic experiment. Cytosolic ribosomal proteins have a substantially higher mean log_2_ LFQ intensity than the detected proteins of mitochondrial ribosome and most of the other ribosome-associated proteins, indicating their enrichment in the analyzed fractions (Figure 4, Supplementary Table 4). The presence of mitochondrial ribosomal proteins in the samples could be attributed to their release from the organelles during the mechanical disruption of the cells. Co-purifying proteins whose homologs were reported to physically associate with ribosomal proteins in *S. cerevisiae* were also identified. These include homolog of the nuclear export factor Arx1, mRNA turnover and ribosome assembly factor Mrt4, elongation factor Tef1, alpha and beta chains of tubulin (Tub1, Tub2), suppressor protein Stm1, alpha subunit of the pyruvate dehydrogenase Pda1 and fatty acid synthases subunits Fas1 and Fas2. Other abundant proteins without known physical interaction with the ribosomes include homologs of NAD-dependent glutamate dehydrogenase Gdh2, clathrin cage assembly protein YAP1801, gamma subunit of translation elongation factor Cam1 and alpha subunit of succinyl-CoA ligase Lsc1 (Supplementary Table 4).

**Fig. 4.**
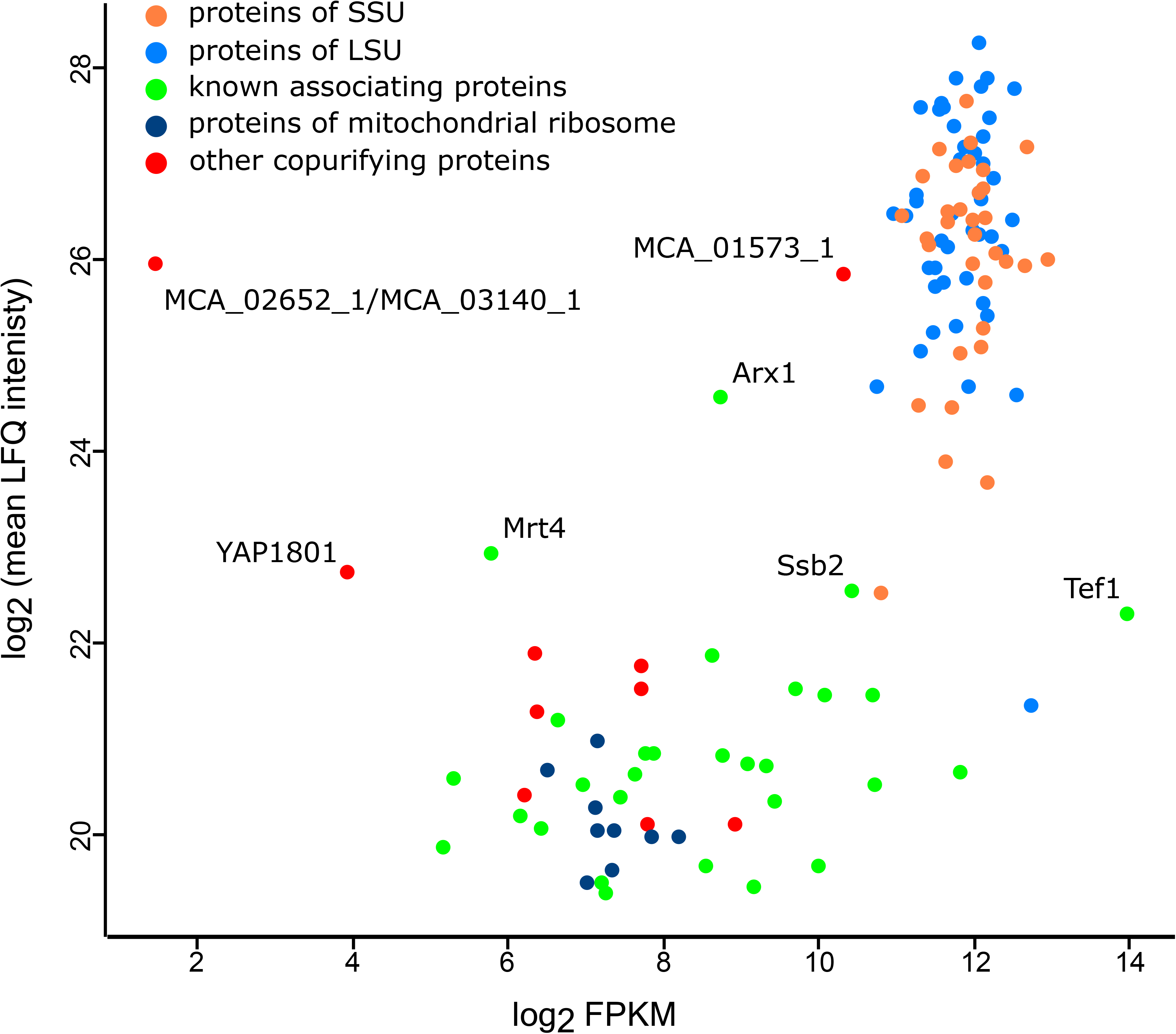
LC-MS/MS analysis of *M. capitatus* cytosolic ribosomes. The results are displayed as a scatter plot of relative quantity (Label Free Quantification LFQ – y-axis) of identified proteins and mRNA coverage (Fragments Per Kilobase per Million mapped fragments FPKM – x-axis) of the gene encoding that particular protein (Brejová et al. 2019a). Among the most abundant non-ribosomal proteins are homologs of Ssa1/Ssa2, Arx1, Mrt4, Ssb2, Tef1, YAP1801, and protein MCA_01573_1.

In addition, several heat-shock proteins of the Hsp70 family were identified in the ribosomal fractions of both *M. capitatus* and *D. albidus* (Supplementary Table 4). In *M. capitatus*, these include the proteins MCA_04550_1/MCA_00692_1, MCA_02011_1, and MCA_03999_1 belonging to SSA, SSB, and SSC subfamily, respectively (for classification of *M. capitatus* Hsp70 proteins see Brejová et al. 2019a). Members of both SSA and SSB subfamilies were reported to play an important role in translation and ribosome biogenesis (Horton et al. 2001, Peisker et al. 2010, Mudholkar et al. 2017, Lee et al. 2021). The SSC homolog presumably represents a contamination by mitochondrial proteins present in the analyzed samples (see above) as this Hsp70 protein associates with mitochondrial protein complexes (Song et al. 2023). The Hsp70 family is expanded in magnusiomycetes (i.e., *M. capitatus* genome contains 47 genes coding for Hsp70 proteins *vs.* 14 and 9 members found in *S. cerevisiae* and *D. albidus*, respectively; Brejová et al. 2019a). This raises a question as to whether the expansion of the Hsp70 family is associated with the reduction of rRNAs. Although we identified MCA_03140_1/MCA_02652_1 classified as novel members of the Hsp70 family (Brejová et al. 2019a) in the ribosomal fractions of *M. capitatus*, the overall expression of the corresponding genes is very low. Moreover, none of the remaining 35 Hsp70 proteins classified as novel were detected in the ribosomal preparations thus questioning the idea that the loss of ESs was compensated by the expansion of this protein family.

Interestingly, we also identified several abundant proteins whose homologs are absent in *S. cerevisiae*. The most interesting is a 532 AAs long protein MCA_01573_1, (Figure 4, Supplementary Table 4), which contains an interferon-related developmental regulator domain (IPR007701). High abundance of this protein in *M. capitatus* ribosomal fraction (log2 mean LFQ intensity 25.85) and also its high expression in the RNA-Seq experiment (FPKM 1267.4, percentile rank 97.0%, Brejová et al. 2019a) indicate that it may represent a *bona fide* ribosomal protein. This conclusion is also supported by the high abundance of this protein in the ribosomal fraction from *D. albidus* (Supplementary Table 4). A homology-based search revealed homologs of MCA_01573_1 in the yeasts of *Dipodascaceae* family: e.g., *M. ingens* (MIA_02712_1) and *G. candidus* (A0A0J9X7V6_GEOCN), as well as in several yeasts from other phylogenetic clades: e.g., *Trichomonascus ciferrii* (A0A642VAI2_9ASCO), *C. albicans* (A0A1D8PDV1_CANAL) and *Blastobotrys adeninivorans* (A0A060T959_BLAAD). A more sensitive remote homology search HHpred (Zimmermann et al. 2018; Gabler et al. 2020) identified two related proteins an interferon-related developmental regulator 1 (IFRD1, E-value 3.1e-57) of the fruit fly *Drosophila melanogaster* and a rabbit IFRD2 (E-value 3.4e-62). They both directly bind to the ribosomes and repress the translation (Brown et al. 2018; Hopes et al. 2022). The binding of IFRD1/IFRD2 proteins to the ribosomes is thought to have a role during the cell differentiation (Brown et al. 2018; Hopes et al. 2022) and in preservation of ribosomes during cellular stress conditions (Brown et al. 2018).

Similarly, two more proteins without homology to *S. cerevisiae* were identified in the ribosomal fractions of *M. capitatus,* namely MCA_02502_1 (log2 mean LFQ intensity 21.29) and MCA_03313_1 (log2 mean LFQ intensity 21.77). These proteins are of lower abundance and expression than *bona fide* ribosomal proteins (Supplementary Table 4). In case of MCA_02502_1, no domain was identified, however, HHpred search revealed a ribosome biogenesis factor Alb1 as a related protein (E-value 3.2e-32). MCA_03313_1 contains Ccdc 124/Oxs1 domain (IPR010422) that can be found in yeast Lso2 ribosome-associated protein (LSO2_YEAST). The similarity with Lso2 protein was further supported by a HHpred search (E-value 8.5e-13). HHpred also identified homology to the human CCDC124 protein (CC124_HUMAN, E-value 5.9e-41) that was shown to associate with 80S ribosomes bound by Nsp1 protein of SARS-CoV-2 (Thoms et al. 2020). The functions of IFRD1/IFRD2 homolog together with MCA_02502_1 and MCA_03313_1 have not been yet characterized in magnusiomycetes and their roles in the process of protein synthesis need to be further addressed experimentally.

### Proteins involved in rRNA biogenesis and ES-associated proteins are conserved in magnusiomycetes

Several reports point to the role of ESs in rRNA biogenesis, maturation and stability (Sweeney et al. 1994; Jeeninga et al. 1997; van Nues et al. 1997; Bradatsch et al. 2012; Ramesh and Woolford 2016; Vos and Kothe 2022). Therefore, we also investigated the genes whose products cleave, process, aid the assembly, modify, or export the rRNA molecules from the nucleus. To identify magnusiomycete orthologs, we used a list of almost 200 proteins (adopted from Woolford and Baserga (2013); Klinge and Woolford (2019) and citations therein) as queries in Blast searches and only the best reciprocal matches with *S. cerevisiae* were included in the final set (Supplementary Table 5). In addition to *M. capitatus*, our analysis also included *M. ingens* (Brejová et al. 2019a), *D. albidus* (Shen et al. 2018), and *G. candidus* (Morel et al. 2015) belonging to the *Dipodascaceae* family. While *M. ingens* lacks the same rRNA segments as *M. capitatus*, *D. albidus,* and *G. candidus* possess canonical yeast rRNAs (Figure 1, Supplementary Figure 2A-D). In the genomes of all four species, we identified homologs of almost all proteins from the list. Homologs of the genes encoding Nop19 and Utp9 were not found in these species and a homolog of YBL028C protein was found only in *D. albidus.* Moreover, we were unable to identify a homolog of putative RNA exonuclease Rex4 in *M. capitatus* and *M. ingens* and a homolog of pre-rRNA processing protein Slx9 was identified only in *G. candidus* (Supplementary Table 5). In addition, several proteins which display low or no homology to *S. cerevisiae* queries were identified by the presence of corresponding PFAM domains. These include the proteins Alb1, Mtr2, Rrt14 and Utp8 (all four species), Bud21, Cgr1 and Rix1 (*M. capitatus*, *M. ingens*), Cms1 and Fyv7 (*M. ingens, G. candidus*), Utp30 (*M. capitatus, D. albidus*), Faf1 (*M. capitatus*), and Loc1 (*G. candidus*) (Supplementary Table 5). Proteins with low homology or those not identified in the analyzed genomes are in many cases relatively small, not exceeding 200 AAs, which complicates their identification.

Recent studies showed that helices 25ES7 and 63ES27 play a major role in anchoring of nascent peptide modifying complexes (Gómez Ramos et al. 2016; Fujii et al. 2018; Knorr et al. 2019,2023; Wild et al. 2020; Shankar et al. 2020; Krauer et al. 2021). These are involved in N-terminus modification (removal of initiator methionine, acetylation, myristoylation), folding and co-translational targeting (Wild et al. 2004; Wegrzyn and Deuerling 2005; Kramer et al. 2009; Gloge et al. 2014; Giglione et al. 2015). Since the ESs are greatly reduced in magnusiomycetes, we searched for protein subunits of nascent-peptide associating complexes. Specifically, we analyzed subunits of methionine aminopeptidase complex (Map1, Map2), N-terminal acetyltransferase complex A and B (Ard1, Nat1, Nat3, Nat5, Mdm20 and Mak3), gene encoding N-myristoyl transferase (Nmt1), subunits of ribosome associated complex (Zuo1, Ssz1, Ssb1/Ssb2, Egd1, Egd2) and a signal recognition particle complex (Srp14, Srp21, Srp54, Srp68, Srp72, Sec65). In all four examined species, we identified all but homologs of Nat5, Srp14 and Srp21 (Supplementary Table 6). These proteins are relatively short, which may hinder their identification by homology-based searches. In the case of Nat5 and Srp14, we were able to find the potential homologs only by searching the corresponding PFAM domains.

In summary, although we identified several minor changes, the overall inventory of proteins involved in rRNA biogenesis and nascent-peptide associating complexes seems to be unaffected in magnusiomycetes, suggesting that their function may be retained independently of the presence of ESs.

### Reduction of ESs is not limited to *Magnusiomyces/Saprochaete* clade

To investigate whether the rRNAs with reduced ESs occur also in other yeast lineages, we searched for rRNA sequences in available genomic assemblies of species from the subphylum Saccharomycotina (Shen et al. 2018) employing Rfam covariance models. In several cases, the genes for rRNAs were missing from the available assemblies. For these species, we used rRNA sequences from the GenBank database (Supplementary Table 2). Results of comparative analysis indicate that canonical (i.e. *S. cerevisiae*-like) rRNAs are the most common in this subphylum and, presumably, they represent an ancestral form. However, similarly to magnusiomycetes, species of the genera *Kodamaea, Komagataella, Metschnikowia, Phaffomyces, Saturnispora,* as well as *Candida sorboxylosa* classified into *Pichiaceae* family, possess rRNAs with reduced ESs (Figure 5, Supplementary Table 2, Supplementary Files 1-4). These species belong to distinct branches on the phylogenetic tree, indicating that the rRNAs alterations occurred independently multiple times during the evolution of this subphylum. The most striking examples represent *C. sorboxylosa* and yeasts from the genus *Komagataella* (e.g. *K. populi*) that lack ∼650 nt of 25S rRNA and more than 210 nt of 18 rRNA. Similarly to magnusiomycete rRNAs, the majority of the missing sequences correspond to 25ES7 and 63ES27. As these changes occur in multiple phylogenetic lineages, the reduction of ESs appears as a common evolutionary trend (Figure 5, Supplementary Table 2, Supplementary File 4).

**Fig. 5.**
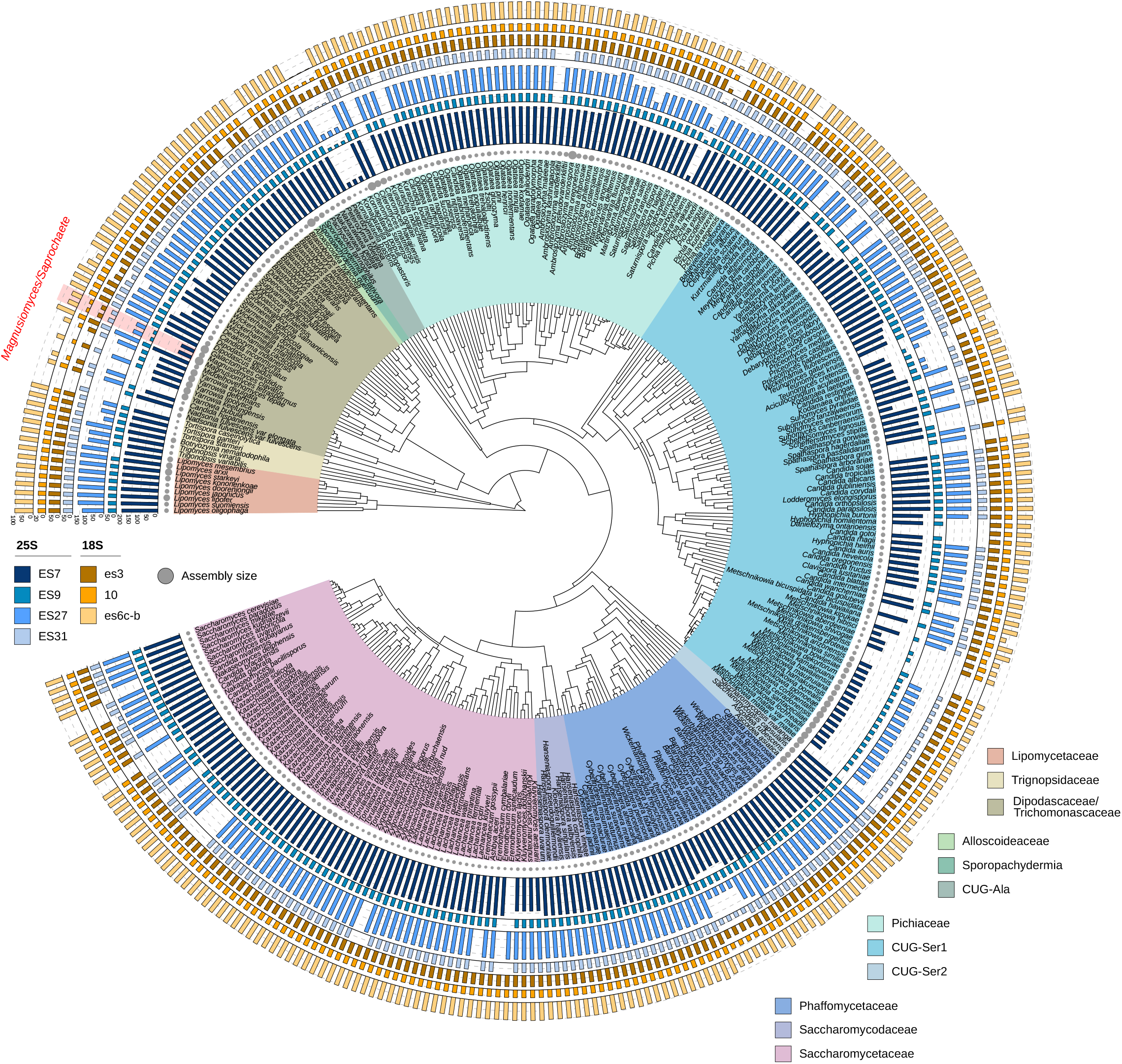
Phylogenetic tree of Saccharomycotina species with sizes of selected rRNA regions. Lengths of segments of SSU stretching over 9es3, helix 10, and 21es6c-es6b (brown colors) and of LSU 25ES7, 31ES9, 63ES27, and 79ES31 (in blue colors) are indicated by bar graphs. In several cases the rRNA sequence is not known/available, resulting in spots with missing data. The sizes of genome assemblies are indicated by gray circles. Evolutionary relationships among the analyzed yeasts are displayed by a phylogenetic tree (adapted from Shen et al. 2018). The clades of subphylum Saccharomycotina are color indicated. For the source data see Supplementary Table 2.

### Evolutionary considerations

Biesiada et al. (2022) have suggested that ESs emerged by a constructive neutral evolution (Stoltzfus 1999; Gray et al. 2010; Muñoz-Gómez et al. 2021), a gradual non-adaptive process that may lead to a high complexity gain of function, and eventually to a dependency. While not functional when acquired, the cells became accustomed to the presence of ESs and employed them as interaction scaffolds, moving from an independent state to dependence (Biesiada et al. 2022). This can be illustrated by ESs in *S. cerevisiae* rRNAs, most of which are essential for cell survival (Ramesh and Woolford 2016). In order for the species to subsequently lose their ribosomal ESs, either a selective pressure has to overcome the acquired functional dependency, or the cellular functions provided by ESs have to become expendable, possibly replaced by other component(s). Both of these scenarios provide an opportunity for the cell to lose ESs and preserve precious resources. In parasitic eukaryotes, the ribosome reduction accompanied the global genome compaction, thus exemplifying a scenario of regressive evolution (Melnikov et al. 2018; Barandun et al. 2019; Hiregange et al. 2022). However, the Saccharomycotina species including magnusiomycetes are free-living organisms and do not exhibit any apparent correlation between the genome size and the length of ESs (Figure 5) and, at the same time, we did not identify compensatory changes in the ribosomal protein inventories or ribosome-interacting partners clearly associated with the shortening of rRNAs. Importantly, we cannot rule out that the truncated ESs retained their biological roles and/or that their loss is compensated by amino acid substitutions in identified proteins. Yet, the dramatic reduction of several ESs found in magnusiomycetes and additional yeast lineages opens up questions regarding their functionality and also points to the importance of further comparative and functional studies of ribosomes in non-model organisms.

## Materials and Methods

### rRNA sequence annotation and phylogenetic analysis

The genome sequences of *M. capitatus*, *M. ingens*, *S. fungicola*, *S. ingens*, and *S. suaveolens* were published previously (Brejová et al. 2019a,b; Hodorová et al. 2019; Lichancová et al. 2019), high contiguity genome assemblies of the remaining magnusiomycete species were determined in our laboratory and will be described elsewhere. The rRNA genes were identified using Rfam 14.0 (Kalvari et al. 2018) and the sequences of 5S, 18S, 25S rRNA, and ITS1 – 5.8S rRNA – ITS2 were deposited in the Genbank database (Supplementary Table 1). The rRNA sequences from *S. cerevisiae* were obtained from GenBank (*RDN18-1* (NR_132213.1), *RDN25-1* (NR_132209.1), *RDN58-1* (NR_132211.1), *RDN5-1* (NR_132215.1), and *RDN37-1* (NR_132207.1); Johnston et al. 1997) and aligned with magnusiomycete counterparts using MAFFT (version 7.388; Katoh and Standley, 2013; Geneious package R11; Biomatters). For phylogenetic analysis, the resulting alignments were concatenated and the columns with more than 50 % gaps were omitted. The phylogenetic tree in Figure 1 was constructed using FastTree (version 2.1.11; Price et al. 2010; Geneious package R11; Biomatters) with the Jukes-Cantor model.

To explore rRNA sequences of other Saccharomycotina species, publicly available genome sequences (Supplementary Table 2) were searched for with an Infernal software package (version 1.1.2, Nawrocki and Eddy 2013). Command *cmsearch* was used to search for genes encoding rRNAs using eukaryotic covariance models of 5S rRNA (RF00001), 5.8S rRNA (RF00002), SSU rRNA (RF01960) and LSU rRNA (RF02543) downloaded from RFAM database v. 14.4 (https://rfam.org/; Kalvari *et al*. 2018). Top scoring results were extracted and manually evaluated. Where possible, missing or short sequences were replaced by sequences from GenBank database and the 5’ and 3’ termini were trimmed to correspond to the RFAM database rRNA annotation (see Supplementary Table 2). Then, all the sequences were aligned to the CM model using *cmalign* command, the sizes of regions spanning ESs were calculated and the data was visualized as bar graphs on the phylogenetic tree of Saccharomycotina species (Shen et al. 2018) using iTOL v. 6.5.8 software (Letunic and Bork 2021).

### Modeling of rRNA structures

The trimmed rRNA sequences were aligned with Clustal Omega (version 1.2.4; Sievers et al. 2011) and further mapped onto the secondary structures of rRNAs from *S. cerevisiae* based on crystallography data (Ben-Shem et al. 2011; Bernier et al. 2014; Petrov et al. 2014b). Secondary structures of rRNA segments that are altered in magnusiomycetes were modeled using Mfold (Zucker, 2003) and adjusted manually. These regions were then highlighted in the 3D structure of *S. cerevisiae* ribosome with modeled 25ES7a and 63ES27a helices (Ben-Shem et al. 2011, RCSB PDB ID: 4V88; Knorr et al. 2019, RCSB PDB ID: 6HD7) using PyMOL (The PyMOL Molecular Graphics System, Version 2.3.0. Schrödinger, LLC.). The secondary structure model of the IC1 intron identified in the 25S rRNA gene of *S. chiloensis* by Rfam searches was built essentially as described in Nawrocki et al. (2018). First, the alignment file containing structural information of available IC1 introns was downloaded from GISSD database (Zhou et al. 2008) (http://www.rna.whu.edu.cn/gissd/; accessed on March 30, 2023). The covariance model was built and calibrated using *cmbuild* and *cmcalibrate*, respectively, and searched against the 25S rRNA gene of *S. chiloensis* with *cmsearch* (Nawrocki and Eddy, 2013). The intron secondary structure was visualized using Varna applet (Darty et al. 2009) and redrawn to follow the conventions (Cech et al. 1994).

### Identification of ribosomal proteins

Homologs encoding ribosomal proteins were identified in the annotated genome sequence of *M. capitatus* by Blast searches (Altschul et al. 1990). In some cases, related proteins were identified using a remote homology search HHpred (Zimmermann et al. 2018, Gabler et al. 2020). Where necessary, original automatic annotations were manually adjusted according to data from an RNA-seq experiment (Brejová et al. 2019a). Due to its small size (25 AA), the gene for the homolog of ribosomal protein eL41 (MCA_10722_1) was identified manually based on conserved synteny with *S. cerevisiae* and high coverage in RNA-seq data. For comparison, *D. albidus* was used as a phylogenetically close yeast with rRNA containing ESs missing in magnusiomycete species (Shen et al. 2018). In this case, only homologs identified by Blast without further correction were used.

### Preparation of cytosolic ribosomes

*M. capitatus* NRRL Y-17686 (CBS 197.35) and *D. albidus* NRRL Y-12859 (CBS 766.85) cells were grown overnight in a liquid YPGal medium (1 % (w/v) yeast extract, 2 % (w/v) peptone, and 2 % (w/v) galactose) till the late exponential phase. Hyphae and yeast cells were collected by filtration. The fractions of cytosolic ribosomes were prepared essentially as described in Ben-Shem et al. (2010,2011) except that the carbon starvation step was omitted. The cells (about 4 g of wet weight) were washed with ice-cold water and with cold Solution A (0.7 M sorbitol, 50 mM KCl, 30 mM HEPES-KOH pH 7.5, 10 mM MgCl_2_, 0.5 mM EDTA, 2 mM dithiothreitol), resuspended in 30 ml of Solution A supplemented with 1 mM phenylmethylsulfonyl fluoride (PMSF) and 0.175x cOmplete™ (Mini, EDTA-free) protease inhibitor cocktail (Roche) in a 50 ml Falcon tube and disrupted by vortexing with an equal volume of glass beads (425-600 μm; Sigma-Aldrich) in ten 20-second cycles with intermittent cooling on ice for 40 seconds. After decantation, the supernatant was cleared by two rounds of centrifugation at 20,000 ×*g* (10 min, 4 °C, JA-20 rotor, Avanti J-26 XP, Beckman Coulter) followed by centrifugation at 31,900 ×*g* (10 min, 4 °C, JA-20 rotor, Avanti J-26 XP, Beckman Coulter). Next, 30 % (w/v) polyethylene glycol (PEG) 20,000 (Carl Roth) was added to the supernatant to a final concentration of 4 % (w/v). The suspension was incubated on ice for 5 min and centrifuged at 20,000 ×*g* (10 min, 4 °C, JA-20 rotor, Avanti J-26 XP, Beckman Coulter). The pellet was re-centrifuged for 5 min and both supernatants were combined into a new tube. 2 M KCl was added to the supernatant to a final concentration of 130 mM, and the suspension was incubated on ice for 5 min. Subsequently, the concentration of PEG 20,000 was adjusted to 9 % (w/v), and the suspension was incubated on ice for 10 min. Precipitated ribosomes were pelleted at 20,000 ×*g* (10 min, 4 °C, JA-20 rotor, Avanti J-26 XP, Beckman Coulter). To remove residual supernatant the pellet was re-centrifuged for 5 min at 20,000 ×*g*. The ribosomal fraction was resuspended in 2 ml of Solution B (0.5 M sorbitol, 150 mM KCl, 30 mM HEPES-KOH pH 7.5, 10 mM MgCl_2_, 2 mM dithiothreitol, 0.5 mM EDTA) containing 1 mM PMSF and 1x cOmplete™, Mini, EDTA-free protease inhibitor cocktail (Roche). The ribosomes were further purified by centrifugation at 59,727 ×*g* (15 hours, 4 °C, SW 32.1 Ti rotor, Optima™ L-100 XP, Beckman Coulter) in a linear (15 – 30 %) sucrose gradient prepared in 20 mM HEPES-KOH pH 7.5, 120 mM KCl, 8.3 mM MgCl_2_, 0.3 mM EDTA, 2 mM dithiothreitol. After the centrifugation, 500 μl fractions were collected starting from the top of the tube, and the absorbance at 260 nm was recorded using a NanoDrop^TM^ One (Thermo Fisher Scientific). 40 μl aliquots from four consecutive fractions were pooled and RNA was extracted using Direct-zol RNA Kit (Zymo Research), analyzed on a 1 % (w/v) agarose gel, and stained using ethidium bromide (0.5 μg/ml) (Supplementary Figure 4).

### Protein identification by LC-MS/MS

Identification of proteins from the ribosomal fractions was done in three biological replicas. Seven fractions with the highest A_260_ containing rRNAs were pooled, precipitated with four volumes of cold acetone (–20 °C), and incubated overnight in a freezer (–20 °C). The precipitate was then pelleted by centrifugation at 16,100 ×*g* (10 min, 4 °C, Eppendorf 5415R). The pellet was washed twice with 80 % (v/v) acetone (–20 °C), air-dried for 30 min at room temperature and suspended in 8 M urea in 50 mM ammonium bicarbonate and 5 mM dithiothreitol. The sample was incubated with occasional shaking for 60 min at 37 °C to dissolve the pellet and reduce disulfide bonds. A sample corresponding to 5 μg of proteins (determined using a Bradford protein assay (Bio-Rad)) was alkylated by 15 mM iodoacetamide for 30 min at room temperature in the dark. Following alkylation, the sample was diluted with 4 volumes of 50 mM ammonium bicarbonate and CaCl_2_ was added to a final concentration of 1 mM. Proteins were digested with modified trypsin (sequencing grade, porcine, Promega) in enzyme:protein ratio of 1:50 overnight at 37 °C. The solution was acidified by the addition of trifluoroacetic acid to a final concentration of 0.5 % (v/v) and loaded into C18 resin containing stage tips. Peptides were washed with 0.1 % (v/v) trifluoroacetic acid, eluted with 2 x 100 μl of 70 % (v/v) acetonitrile in 0.5 % (v/v) trifluoroacetic acid into low-binding microcentrifugation tubes (Protein LoBind, Eppendorf), and vacuum-dried in Vacufuge Concentrator Plus (Eppendorf). Peptides were solubilized in 7 μl of 2 % (v/v) acetonitrile in 0.5 % (v/v) trifluoroacetic acid and analyzed by LC-MS/MS using an Orbitrap Elite mass spectrometer (Michalski et al. 2012) in two technical replicas. Raw data were analyzed using MaxQuant (version 2.1.3.0) (Cox and Mann 2008). Standard configuration was used for MS spectra, the maximum number of modifications per peptide was set to 3 and label-free quantification was turned on. The MS/MS spectra were searched against predicted *M. capitatus* (Brejová et al. 2019a) and *D. albidus* (Shen et al. 2018) proteomes. Peptides and corresponding proteins were analyzed and processed with Perseus (version 2.0.6.0) (Tyanova et al. 2016). Contaminating proteins, reverse proteins, proteins identified only by site, and proteins not identified in all three biological replicas were removed from the final dataset. In case of *M. capitatus*, the mean values of LFQ intensities from biological replicas were log_2_ transformed and plotted against FPKM count from the RNA-seq analysis (Brejová et al. 2019a). To functionally annotate the identified proteins, *S. cerevisiae* was used as a reference and BlastP (Altschul et al. 1990) searches were performed to identify homologous proteins.

## Data Availability

The sequences of magnusiomycete rRNAs were deposited in the GenBank database under the accession numbers listed in Supplementary Table 1. Genome assemblies were retrieved from the publicly available databases (Supplementary Table 2). The mass spectrometry proteomics data have been deposited to the ProteomeXchange Consortium via the PRIDE (Perez-Riverol et al. 2022) partner repository with the dataset identifier PXD043413.

## Supplementary Material

Supplementary data are available online.

## Acknowledgements

We would like to thank Cletus P. Kurtzman and James Swezey (Agricultural Research Service, Peoria, IL, USA) for providing us with the strains of *M. capitatus* and *D. albidus*. We also thank Viktória Hodorová and Hana Lichancová (Comenius University in Bratislava, Slovakia) for sharing their unpublished sequence data with us. This research was supported by grants from the Slovak Research and Development Agency (APVV 18-0239 and APVV 22-0144 (to J.N.), APVV 19-0068 (to L.T.)), and the Slovak Grant Agency (VEGA 1/0061/20 (to L.T.), VEGA 1/0463/20 (to B.B.), VEGA 1/0538/22 (to T.V.), VEGA 1/0234/23 (to J.N.)). Additional support was provided by the Advancing University Capacity and Competence in Research, Development and Innovation (ACCORD) project.

## Supplementary Materials

**Supplementary Table 1** – Yeast strains and rRNAs accession numbers in the GenBank database.

**Supplementary Table 2** – List of genome assemblies and rRNA source data.

**Supplementary Table 3** – Inventory of ribosomal proteins in *M. capitatus* and *D. albidus*.

**Supplementary Table 4** – Proteins identified by LC-MS/MS ribosomal fractions of *M. capitatus* and *D. albidus*.

**Supplementary Table 5** – List of proteins involved in rRNA biogenesis.

**Supplementary Table 6** – List of proteins associating with nascent peptide.

**Supplementary Figure 1.** – 2D model of the Group I intron present in *S. chiloensis* 25S rRNA gene. The structure was built using a covariance model of known IC1 introns and redrawn to fulfill the convention by Cech et al. (1994). The intronic sequence is indicated by capital letters and flanking exon sequence of 25S rRNA by lower case letters. The red arrows point at the 5’ and 3’ splice sites, respectively. The 5’ to 3’ direction is shown with black arrowheads. The canonical Watson-Crick pairing is indicated with solid lines, G○U wobble base pairs with empty circles and non-Watson-Crick pairing with solid circles. Base pairs predicted by the covariance model are in black and manually altered bonds are in orange. The sequence of the intron is 437 nt long. The model fits a common structure of IC1 introns very well. Compared to the *Tetrahymena thermophila* intron (Cech et al. 1994), the P2.1 of *S. chiloensis* intron is expanded due to the addition of 39 nt and the P7 helix appears to be reduced.

**Supplementary Figure 2.** – Sequence alignments of **(A)** 5S, **(B)** 18S, **(C)** 5.8S, and **(D)** 25S rRNAs. The sequences were trimmed according to Rfam annotation and aligned with Clustal Omega program. rRNA helices numbering was adopted from Petrov et al. (2014b) and Bernier et al. (2014) and displayed above the alignments. The species abbreviations are: *S. cerevisiae* (SacCer), *S. psychrophila* (SapPsy), *D. albidus* (DipAlb), *G. candidus* (GalCan), *M. capitatus* (MagCap), *M. spicifer* (MagSpi), *M. clavatus* (MagCla), *M. magnusii* (MagMag), *S. gigas* (SapGig), *S. suaveolens* (SapSua), *M. tetrasperma* (MagTet), *S. fungicola* (SapFun), *S. saccharophila* (SapSac), *M. starmeri* (MagSta), *S. quercus* (SapQue), *M. ovetensis* (MagOve), *S. chiloensis* (SapChi), *M. ingens* (MagIng), and *S. ingens* (SapIng).

**Supplementary Figure 3.** – Secondary structures of *M. capitatus* rRNAs. **(A)** 18S rRNA, **(B)** 5S, 5.8S, and 25S rRNAs. *S. cerevisiae* rRNAs were used as templates; segments missing in *M. capitatus* are indicated by dotted red lines. Numbering of helices was adapted from Petrov *et al*. (2014b) and Bernier *et al*. (2014).

**Supplementary Figure 4.** – Preparation of ribosomal fractions from *M. capitatus* **(A)** and *D. albidus* **(B)**. The crude ribosomes were further separated by centrifugation in a sucrose gradient, fractions were collected starting from the top of the centrifugation tube. The A_260_ was measured by a spectrophotometer (left), RNA samples were extracted from aliquots pooled from four consecutive fractions, and analyzed on agarose gel (right). Pooled fractions in each aliquot are indicated above the gel. A representative ribosome isolation experiment and fraction analysis out of the three biological replicas is shown for each organism.

Supplementary File 1 – Yeast 5S rRNA sequences aligned to a covariance model (Stockholm format).

Supplementary File 2. – Yeast 5.8S rRNA sequences aligned to a covariance model (Stockholm format).

Supplementary File 3. – Yeast 18S rRNA sequences aligned to a covariance model (Stockholm format).

Supplementary File 4. – Yeast 25S rRNA sequences aligned to a covariance model (Stockholm format).

Supplementary Movie 1. – A model of the cytosolic ribosome of *M. capitatus.* The colors are the same as in Figure 3. The movie was produced in PyMOL (The PyMOL Molecular Graphics System, Version 2.3.0. Schrödinger, LLC.).

