## Supplementary_Figures_1-4 for "Reduction of ribosomal expansion segments in yeast species of the *Magnusiomyces/Saprochaete* clade"

**Supplementary Figure 4. -** Preparation of ribosomal fractions from *M. capitatus* (**A**) and *D. albidus* (**B**). The crude ribosomes were further separated by centrifugation in a sucrose gradient, fractions were collected starting from the top of the centrifugation tube. The $A_{260}$ was measured by a spectrophotometer (left), RNA samples were extracted from aliquots pooled from four consecutive fractions, and analyzed on agarose gel (right). Pooled fractions in each aliquot are indicated above the gel. A representative ribosome isolation experiment and fraction analysis out of the three biological replicas is shown for each organism.

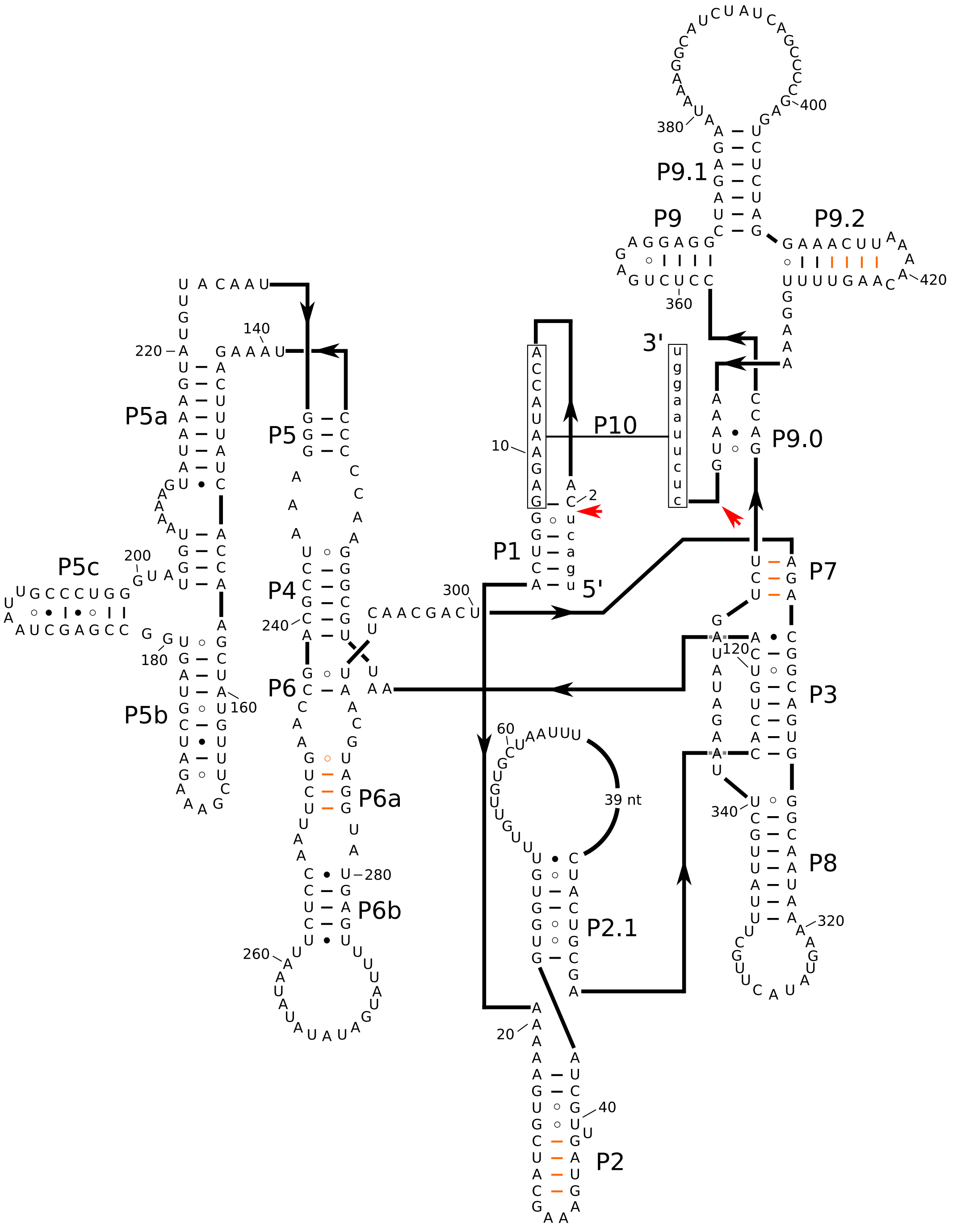

P1
P2
P2.1
P3
P4
P5
P5a
P5b
P5c
P6
P6a
P6b
P7
P8
P9
P9.0
P9.1
P9.2
P10
5'
3'
39 nt

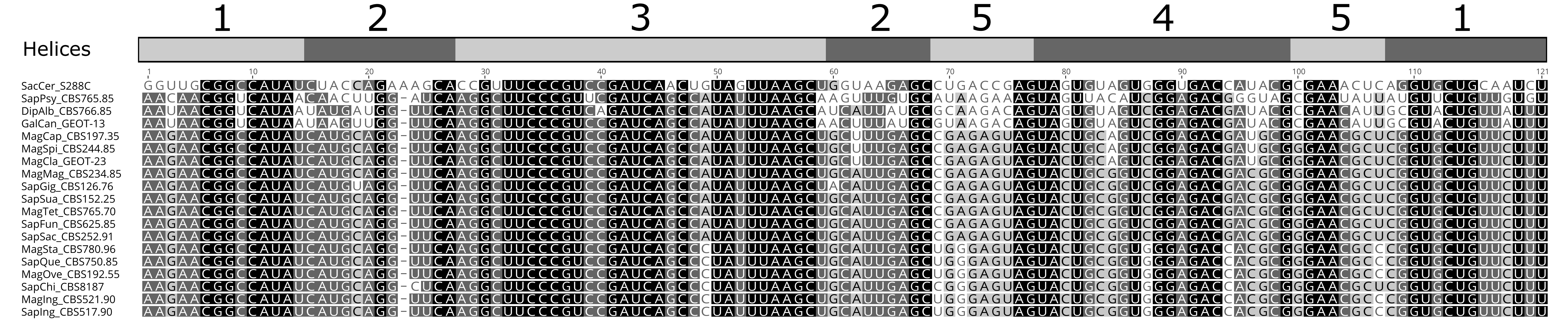
1
2
3
2
5
4
5
1
Helices
1
10
20
30
40
50
60
70
80
90
100
110
121
SacCer_S288C
SapPsy_CBS765.85
DipAlb_CBS766.85
GalCan_GEOT-13
MagCap_CBS197.35
MagSpi_CBS244.85
MagCla_GEOT-23
MagMag_CBS234.85
SapGig_CBS126.76
SapSua_CBS152.25
MagTet_CBS765.70
SapFun_CBS625.85
SapSac_CBS252.91
MagSta_CBS780.96
SapQue_CBS750.85
MagOve_CBS192.55
SapChi_CBS8187
MagIng_CBS521.90
SapIng_CBS517.90

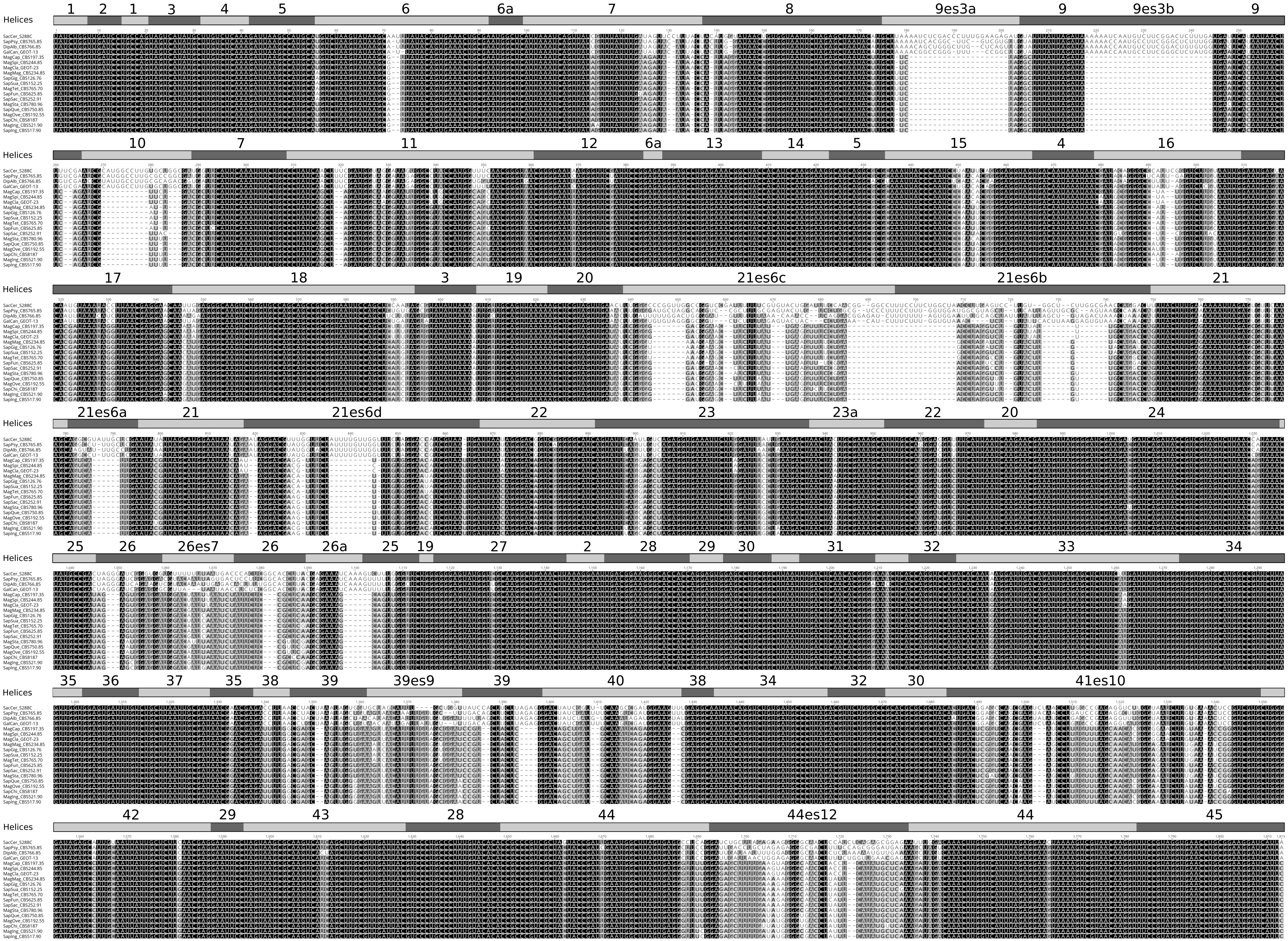

Helices
1 2 1 3 4 5 6 6a 7 8 9es3a 9 9es3b 9
Helices
10 7 11 12 6a 13 14 5 15 4 16
Helices
17 18 3 19 20 21es6c 21es6b 21
Helices
21es6a 21 21es6d 22 23 23a 22 20 24
Helices
25 26 26es7 26 26a 25 19 27 2 28 29 30 31 32 33 34
Helices
35 36 37 35 38 39 39es9 39 40 38 34 32 30 41es10
Helices
42 29 43 28 44 44es12 44 45

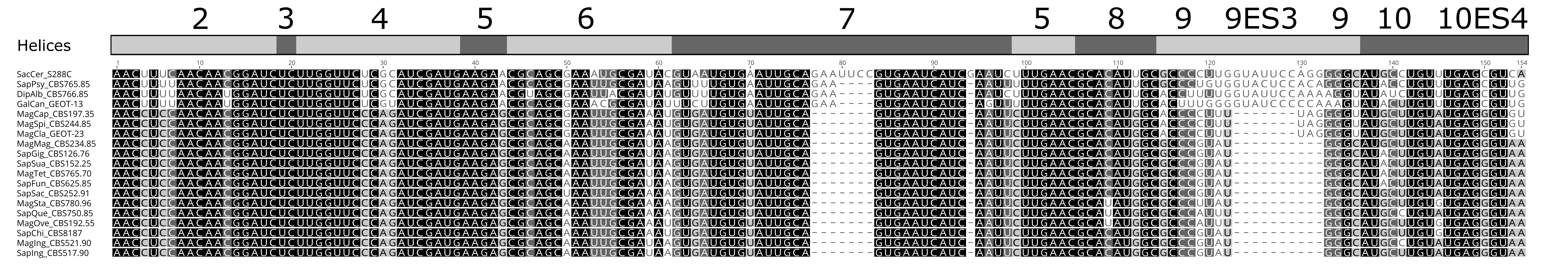
2
3
4
5
6
7
5
8
9
9ES3
9
10
10ES4
Helices
SacCer_S288C
SapPsy_CBS765.85
DipAlb_CBS766.85
GalCan_GEOT-13
MagCap_CBS197.35
MagSpi_CBS244.85
MagCla_GEOT-23
MagMag_CBS234.85
SapGig_CBS126.76
SapSua_CBS152.25
MagTet_CBS765.70
SapFun_CBS625.85
SapSac_CBS252.91
MagSta_CBS780.96
SapQue_CBS750.85
MagOve_CBS192.55
SapChi_CBS8187
MagIng_CBS521.90
SapIng_CBS517.90

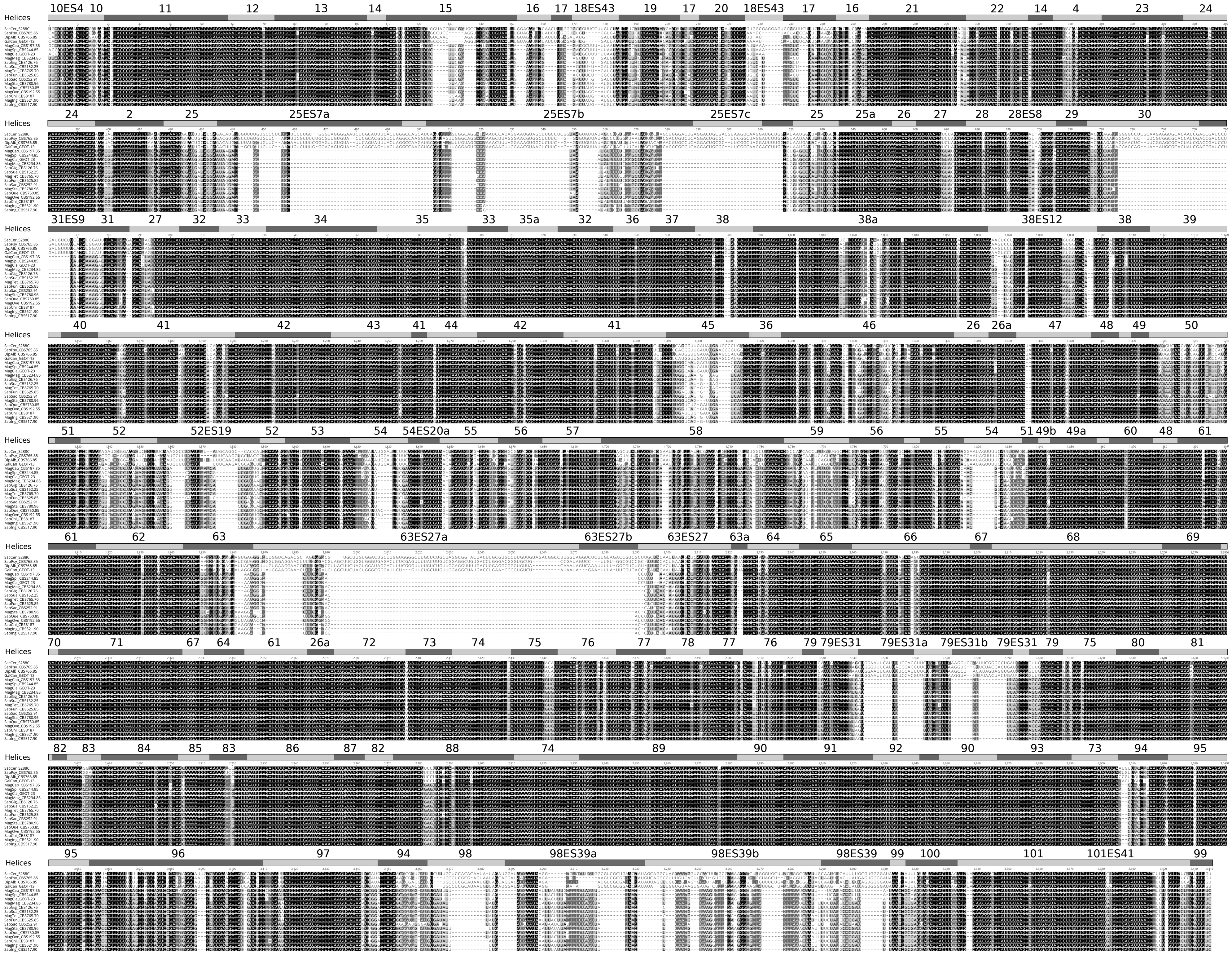
Helices
10ES4 10 11 12 13 14 15 16 17 18ES43 19 17 20 18ES43 17 16 21 22 14 4 23 24
24 2 25 25ES7a 25ES7b 25ES7c 25 25a 26 27 28 28ES8 29 30
31ES9 31 27 32 33 34 35 33 35a 32 36 37 38 38a 38ES12 38 39
40 41 42 43 41 44 42 41 45 36 46 26 26a 47 48 49 50
51 52 52ES19 52 53 54 54ES20a 55 56 57 58 59 56 55 54 51 49b 49a 60 48 61
61 62 63 63ES27a 63ES27b 63ES27 63a 64 65 66 67 68 69
70 71 67 64 61 26a 72 73 74 75 76 77 78 77 76 79 79ES31 79ES31a 79ES31b 79ES31 79 75 80 81
82 83 84 85 83 86 87 82 88 74 89 90 91 92 90 93 73 94 95
95 96 97 94 98 98ES39a 98ES39b 98ES39 99 100 101 101ES41 99

# 18S

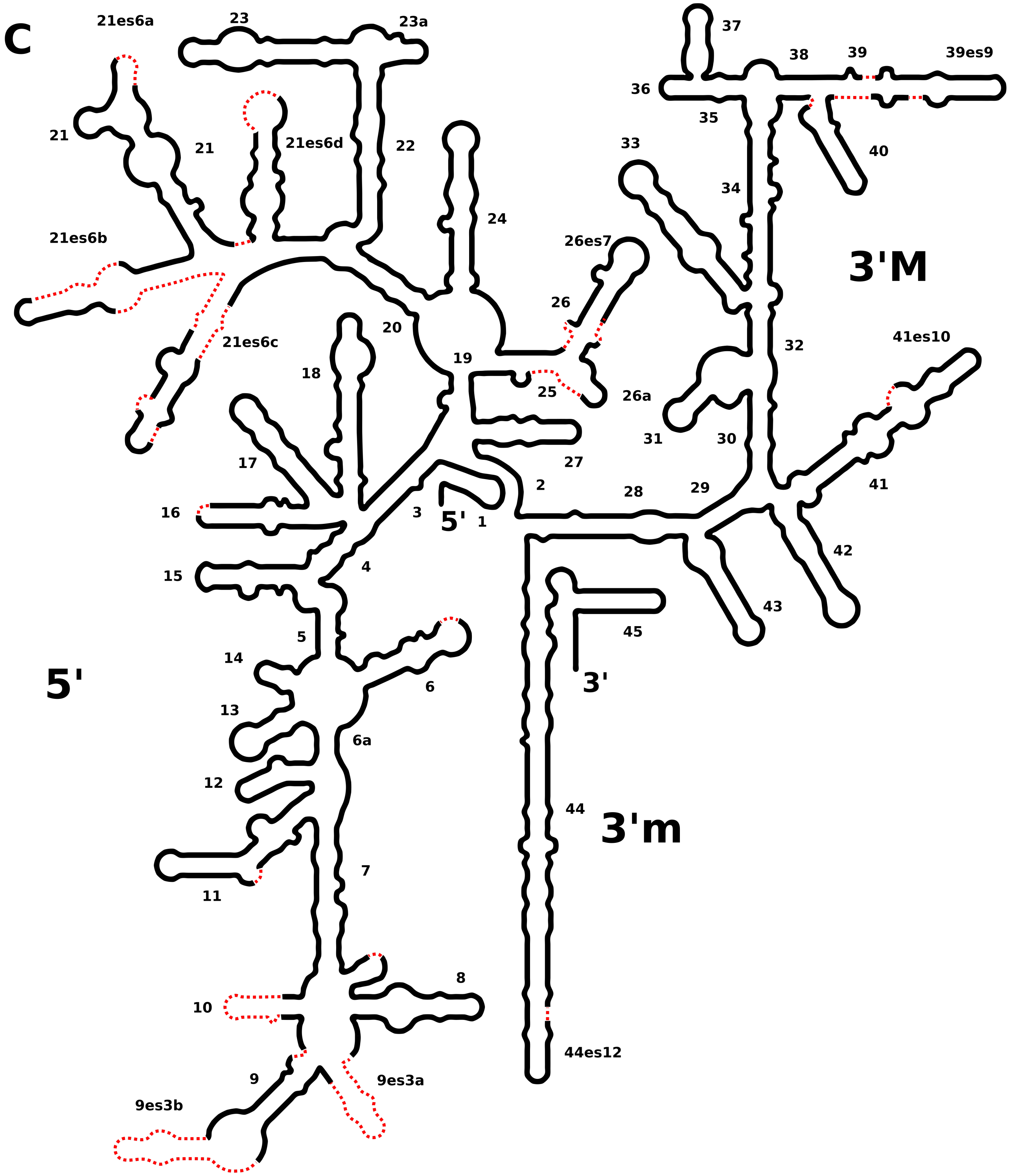

C
21es6a
23
23a
37
38
39
39es9
36
35
21
21
21es6d
22
33
40
34
24
26es7
21es6b
3'M
26
20
21es6c
19
32
41es10
18
25
26a
31
30
17
27
2
28
29
41
16
3
5'
1
15
4
42
43
45
5
14
6
5'
3'
13
6a
12
44
3'm
11
7
8
10
44es12
9
9es3a
9es3b

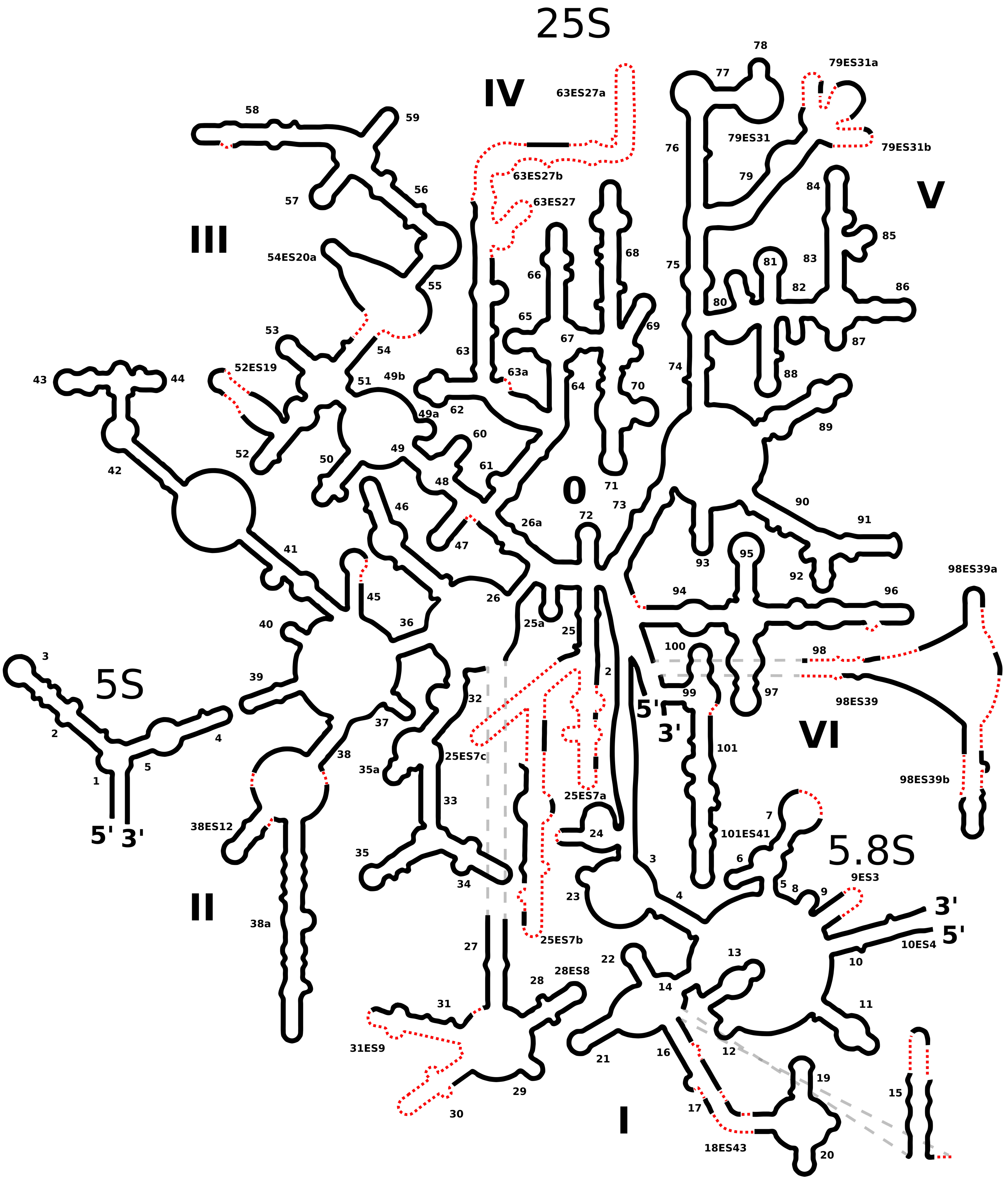

25S
IV
63ES27a
58
59
78
77
79ES31a
76
79ES31
79ES31b
79
63ES27b
63ES27
56
57
III
84
V
85
68
54ES20a
75
81
83
66
82
86
80
55
65
69
53
67
87
54
63
63a
74
88
43
52ES19
44
51
49b
64
70
62
49a
60
89
52
42
50
49
61
48
71
0
73
90
46
72
26a
91
47
41
93
95
92
98ES39a
45
94
96
26
40
36
25a
25
100
98
3
39
5S
2
99
97
98ES39
32
5'
3'
37
2
4
101
VI
38
25ES7c
35a
5
1
98ES39b
25ES7a
33
7
5'
3'
38ES12
24
101ES41
5.8S
35
6
3
9ES3
34
23
4
5
8
9
3'
II
38a
5'
25ES7b
10ES4
27
10
22
13
28ES8
28
14
31
11
21
16
12
31ES9
29
19
15
30
I
17
18ES43
20

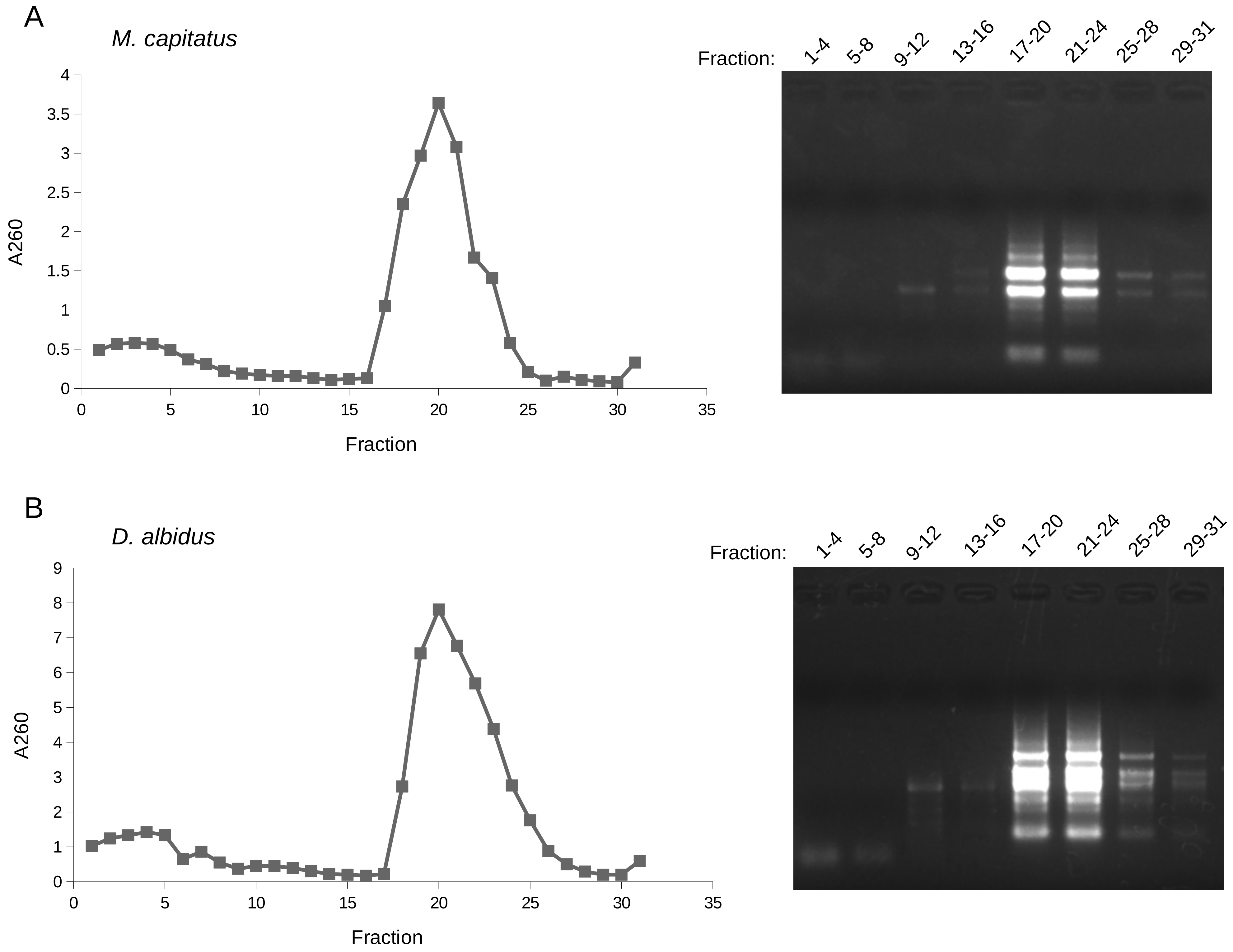
A
M. capitatus
A260
4
3.5
3
2.5
2
1.5
1
0.5
0
0
5
10
15
20
25
30
35
Fraction
Fraction:
1-4
5-8
9-12
13-16
17-20
21-24
25-28
29-31
B
D. albidus
A260
9
8
7
6
5
4
3
2
1
0
0
5
10
15
20
25
30
35
Fraction
Fraction:
1-4
5-8
9-12
13-16
17-20
21-24
25-28
29-31
